# Spectral theory of stochastic gene expression: a Hilbert space framework

**DOI:** 10.1101/2025.05.19.654994

**Authors:** Bingjie Wu, Ramon Grima, Chen Jia

## Abstract

A survey of the literature reveals notable discrepancies among the purported exact results for the spectra of stochastic gene expression models. For self-repressing gene circuits, previous studies ([Phys. Rev. Lett. 99, 108103 (2007)], [Phys. Rev. E 83,062902 (2011)], [J. Chem. Phys. 160, 074105 (2024)], and [bioRxiv 2025.02.05.635946 (2025)]) have provided different exact solutions for the eigenvalues of the generator matrix. In this work, we propose a unified Hilbert space framework for the spectral theory of stochastic gene expression. Based on this framework, we analytically derive the spectra for models of constitutive, bursty, and autoregulated gene expression. The eigenvalues and eigenvectors obtained are then used to construct an exact spectral representation of the time-dependent distribution of gene product numbers. The spectral gap between the zero eigenvalue and the first nonzero eigenvalue, which reflects the relaxation rate of the system towards its steady state, is then compared with the prediction of the deterministic model, and we find that deterministic modeling fails to capture the relaxation rate when autoregulation is strong. In particular, our results demonstrate that for infinite-dimensional operators such as in stochastic gene expression models, many conclusions in linear algebra do not apply, and one must rely on the modern theory of functional analysis.

## 1 Introduction

Over the past few decades, significant progress has been made to derive a spectral theory of stochastic gene expression. Since the mRNA and protein numbers can take any nonnegative integer values, the state space of a stochastic gene expression model is usually infinite and the corresponding generator matrix of the Markovian dynamics is usually an infinite-dimensional operator. The spectral theory is crucial for the study of stochastic gene expression dynamics, mainly for the following two reasons. First, it is well known that the spectral gap between the zero eigenvalue and the first nonzero eigenvalue of the generator matrix characterizes the relaxation rate of the system towards its steady state [1]. The relaxation rate reflects the ability of gene expression levels to return to equilibrium after environmental changes or signal perturbations, serving as a key indicator of robustness and adaptation of the system. Second, the spectral decomposition method has been widely used to obtain the exact time-dependent solution for stochastic gene expression models [2–4], with different spectral components corresponding to different time scales of the system’s dynamical evolution. Therefore, the spectral theory can reveal the detailed dynamical behavior of the system across different time scales.

Within the continuous-time Markovian framework, some studies model the protein dynamics explicitly and model the gene dynamics implicitly [5, 6]. For such models, the spectral theory has been well developed for both unregulated and autoregulated genes when protein synthesis is constitutive [1], i.e. the molecules are produced one at a time. For unregulated genes, the eigenvalues are discrete and are given by

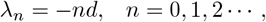

where *d* is the rate of protein degradation. By analyzing the properties of the first nonzero eigenvalue, it has been shown that positive autoregulation always slows down the relaxation kinetics to the steady state, while negative autoregulation always speeds it up [1]. In addition, the time-dependent solution for an autoregulatory feedback loop has been obtained when all eigenvalues of the Markovian model are known [7, 8].

Other studies have considered more realistic models where both the gene and protein dynamics are modeled explicitly [9–14]. For such models, the spectral theory has also been examined for both unregulated and autoregulated genes [3, 4, 15–17]. Since now the gene switching dynamics is taken into account, for unregulated genes, there are two sets of eigenvalues [4]:

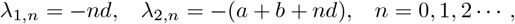

where *d* is the rate of protein degradation, and *a* and *b* are the switching rates between the active and inactive gene states. However, for autoregulatory gene circuits, results on the spectral theory do not agree with each other — previous studies have obtained different purported exact solutions of the eigenvalues. Ramos and coworkers [3, 15–17] claimed that the eigenvalues for self-repressing genes are all real numbers. However, our previous study [4] showed that the eigenvalues for the same system are the roots of a non-trivial continued fraction equation and are not always real as previously claimed. Thus far, there is a lack of a unified mathematical framework for the spectral theory of stochastic gene expression models that can be used to clarify which eigenvalues are correct and which are incorrect.

In the present work, we propose a rigorous Hilbert space framework for the spectral theory of stochastic gene expression. Based on this framework, we analytically derive the spectra for three classical models, including models of constitutive genes, bursty genes, and self-repressing genes (Fig. 1). For constitutive genes, we consider a simple model with only constitutive mRNA synthesis and mRNA degradation. For bursty genes, we consider a model with bursty protein synthesis and protein degradation, where the burst size of protein is assumed to have a geometric distribution, supported by experiments [18]. For self-repressing genes, we consider a more complicated model that includes protein synthesis, protein degradation, gene state switching, as well as autoregulation, whereby the protein produced from a gene represses its own transcription. Our results show that for self-repressing genes, the eigenvalues given in [4] are correct and can be complex numbers. This is contradictory to the real eigenvalues reported in [3, 15–17]. In particular, we demonstrate that for infinite-dimensional operators such as in stochastic gene expression models, many conclusions in linear algebra do not apply, and one must rely on the modern theory of functional analysis.

**Figure 1.**
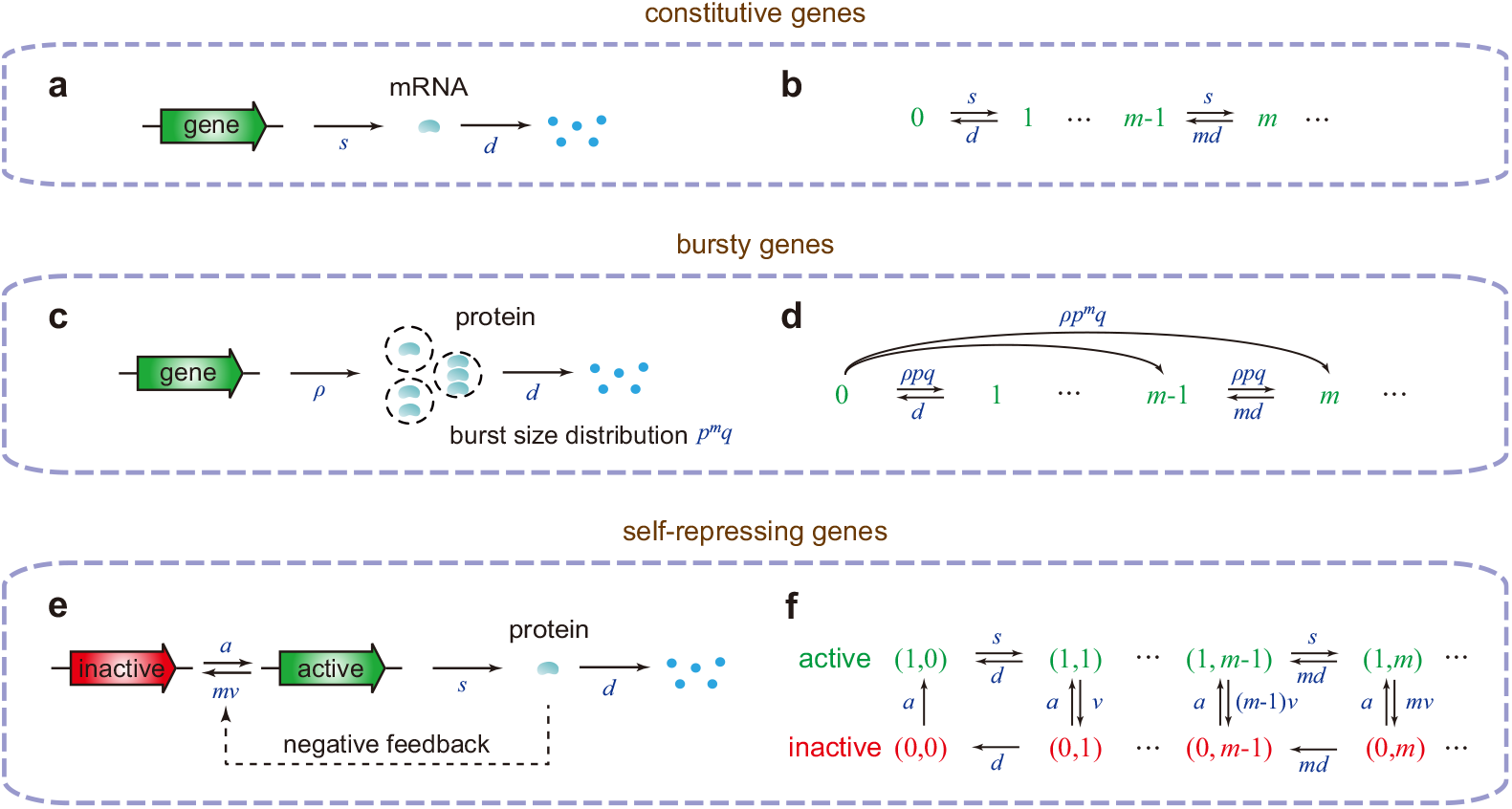
Models. **(a)** Model with constitutive mRNA synthesis and mRNA degradation. **(b)** Transition diagram of the Markovian dynamics for the model shown in (a). **(c)** Model with bursty protein synthesis and protein degradation. **(d)** Transition diagram of the Markovian dynamics for the model shown in (c). **(e)** Model of a self-repressing gene, where the protein expressed from a gene represses its own transcription. **(f)** Transition diagram of the Markovian dynamics for the model shown in (e). In (b) and (d), the microstate of the gene is only described by the gene product number *m* = 0, 1, 2, *· · ·*, while in (f) the microstate state of the gene is described by (*i, m*), where *i* = 0, 1 represents the gene state and *m* = 0, 1, 2, *· · ·* represents the protein number.

As an application of our analytical results, we use the eigenvalues and eigenvectors obtained to construct an exact spectral representation of the time-dependent distribution of gene product numbers for the three models. The magnitude of the real part of the first nonzero eigenvalue, which reflects the relaxation rate to the steady state, is also compared with the prediction of the deterministic model. We demonstrate that deterministic modeling accurately capture the relaxation rate for constitutive and bursty genes, while it fails to reproduce the relaxation rate for self-repressing genes.

## 2 Hilbert space framework

Stochastic gene expression dynamics is typically modeled as a Markov jump process (continuous-time Markov chain) with a discrete state space. Let *V* be the state space of the Markovian model and let *Q* = (*q*_*xy*_)_*x,y*∈*V*_ be the associated generator matrix. The aim of the present paper is to find all the eigenvalues of *Q* for some important gene expression models. If the state space *V* is finite, then all the eigenvalues of the finite-dimensional *Q* can be determined straightforwardly using the standard linear algebraic method — they are simply all the roots of the characteristic equation det(*λI* − *Q*) = 0. However, the numbers of mRNA and protein in living cells can take any nonnegative integer values and hence the state space *V* is usually infinite. In such cases, the eigenvalue problem becomes fundamentally different from the finite-dimensional scenario, and continuing to use the conventional linear algebra method may result in non-physical outcomes.

To resolve this problem, we revisit modern functional analysis theory, where the eigenvalues and eigenvectors of an infinite-dimensional operator must be defined within a given Hilbert or Banach space. For Markov jump processes with infinite state spaces, a common choice for is the Hilbert space [1, 19]:

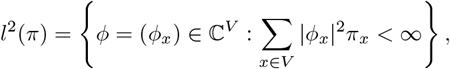

where *π* = (*π*_*x*_)_*x*∈*V*_ is the steady-state distribution of the Markovian model. Note that the generator matrix of a stochastic gene expression model is typically an unbounded operator on *l*^2^(*π*) (this can be readily observed in subsequent sections — the eigenvalues of stochastic gene expression models often have magnitudes that tend to infinity, whereas the eigenvalues of a bounded operator must be bounded by its operator norm [20]). Rigorously, the generator matrix can be viewed as the unbounded operator *Q* : *D*(*Q*) → *l*^2^(*π*) on *l*^2^(*π*) defined by

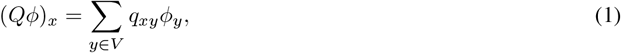

where

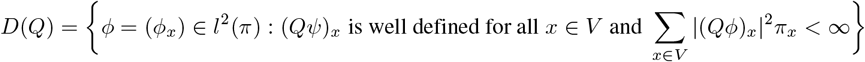

is the domain of the definition of *Q* (for unbounded operators, the domain of definition is only a subspace of *l*^2^(*π*) rather than the entire Hilbert space).

We next recall the definitions of the spectrum and eigenvalues of the generator matrix *Q* [20].

**Definition 1**. The resolvent set of the generator matrix Q is the set of all *λ* ∈ ℂ such that *Q* − *λI* is a one-to-one mapping of *D*(*Q*) onto *l*^2^(*π*) whose inverse belongs to ℬ (*l*^2^(*π*)), where *I* denotes the identity operator and ℬ (*l*^2^(*π*)) denotes the space of all bounded linear operators on *l*^2^(*π*). In other words, *Q* − *λI* should have inverse *S* ∈ ℬ (*l*^2^(*π*)) which satisfies

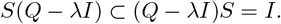

The spectrum of *Q* is the complement of resolvent set of *Q*, i.e. the set of all *λ* ∈ C for which *Q* − *λI* does not have an inverse that is a bounded linear operator.

**Definition 2**. If there exists a nonzero complex-valued column vector *ϕ* = (*ϕ*_*x*_)_*x*∈*V*_ ∈ *l*^2^(*π*) such that *Qϕ* = *λϕ*, then we say that *λ* ∈ ℂ is an eigenvalue of the generator matrix *Q* and *ϕ* is the associated eigenvector. In particular, the set of all eigenvalues *λ* ∈ ℂ is called the point spectrum of *Q*.

A well known result in functional analysis is that for an unbounded operator, the set of eigenvalues, i.e. the point spectrum, must be included in the spectrum, but the spectrum is not just the set of eigenvalues — the spectrum can be decomposed into the point spectrum, continuous spectrum, and residual spectrum [20]. In the present work, we only focus on the point spectrum for stochastic gene expression models.

We emphasize that the nonzero column vector *ϕ* = (*ϕ*_*x*_)_*x*∈*V*_ in Definition 2 corresponds to the right eigenvector of the generator matrix *Q*, and the associated *λ* ∈ ℂ corresponds to the right eigenvalue. Similarly, one can also define the left eigenvalues and eigenvectors of *Q*, where the Hilbert space should be chosen as

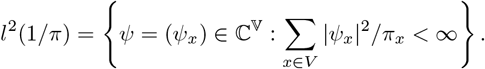

Note that *l*^2^(1*/π*) can be viewed as the dual space of *l*^2^(*π*) [20] in the sense that every *ψ* ∈ *l*^2^(1*/π*) induces a bounded linear functional *f*_*ψ*_ on *l*^2^(*π*) defined by *f*_*ψ*_(*ϕ*) = *x*∈*V ϕ*_*x*_*ψ*_*x*_.

**Definition 3**. If there exists a nonzero complex-valued row vector *ψ* = (*ψ*_*x*_)_*x*∈*V*_ ∈ *l*^2^(1*/π*) such that *ψQ* = *λψ*, then we say that *λ* ∈ ℂ is a left eigenvalue of the generator matrix *Q* and *ψ* is the associated left eigenvector.

For stochastic gene expression models, the left eigenvalues (defined in *l*^2^(1*/π*)) and the right eigenvalues (defined in *l*^2^(*π*)) are generally identical. This equivalence is far from obvious will be discussed in detail in a forthcoming paper. In what follows, we mainly focus on the the left eigenvalues and eigenvectors since they are more amenable to analytical calculation. Note that the above framework is similar to that of quantum mechanics, where any physically meaningful observable must be a Hermitian operator on a Hilbert space and its eigenvalues correspond to all possible outcomes of a measurement of the given observable [21].

## 3 A simple model of constitutive mRNA production

Let *G* denote the gene of interest and let *M* denote the corresponding mRNA. We first focus on a simple gene expression model with constitutive mRNA synthesis (mRNA molecules are produced one at a time) and mRNA degradation. The expression dynamics for a constitutive gene is described by the reactions (Fig. 1(a))

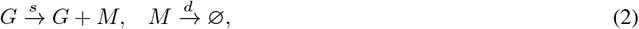

where *s* is the rate of mRNA synthesis and *d* is the rate of mRNA degradation. The microstate of the gene is described by the mRNA number *m* = 0, 1, 2, · · · . The stochastic evolution of the mRNA number in a single cell is characterized by the birth and death process shown in Fig. 1(b). Recall that it is a Markov jump process with generator matrix

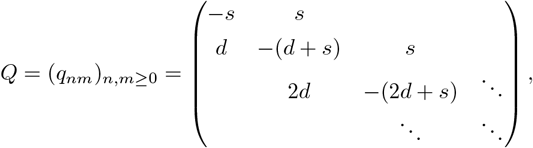

where *q*_*mn*_ denotes the transition rate from microstate *m* to microstate *n*. In steady state, it is well known that the transcript number for the constitutive gene expression model shown in Fig. 1(a) has the Poisson distribution

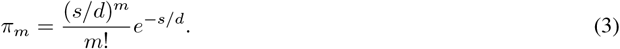

A well-known result is that for this model, the eigenvalues of the generator matrix *Q* are given by [1, 17]

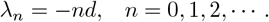

In a recent paper [17], a linear algebraic method was used to compute the eigenvalues for stochastic gene expression dynamics. The logic of the linear algebraic method is as follows. Let *Q* be the generator matrix of a stochastic gene expression model; if one can find a square matrix Φ and a diagonal matrix Λ such that Φ*Q* = ΛΦ, then the diagonal elements of Λ are all the eigenvalues of *Q*. In fact, it can be proved that for any diagonal matrix Λ = diag(*λ*_0_, *λ*_1_, · · ·), one can always find a square matrix Φ = (*ϕ*_*nm*_)_*n,m*≥0_ such that

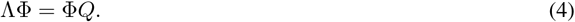

Note that Eq. (4) can be written in component form as

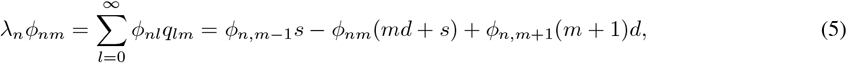

where *ϕ*_*n*,−1_ = 0 by default. Directly deriving an explicit expression of *ϕ*_*nm*_ from the above recurrence relation is difficult and hence we use the generating function method. To proceed, we introduce the generating functions

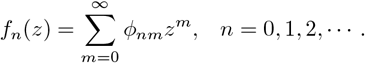

Thus, Eq. (5) can be expressed as the following differential equation:

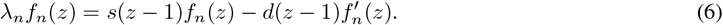

For convenience, we set *ϕ*_*n*0_ = 1, which ensures that *f*_*n*_(*z*) = 1. It then follows that Eq. (6) can be solved explicitly as

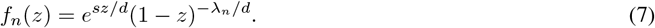

Hence the components of the matrix Φ can be computed in closed form as

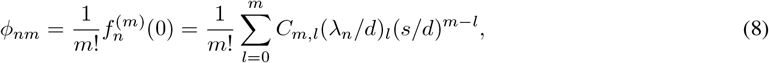

where (*x*)_*l*_ = *x*(*x* + 1) · · · (*x* + *l* − 1) denotes the Pochhammer symbol and *C*_*m,l*_ = *m*! */l*! (*m* − *l*)! denotes the combinatorial number. This implies that, regardless of how the diagonal matrix Λ is chosen, one can always find a matrix Φ such that Eq. (4) holds. Hence, even if Eq. (4) holds, we cannot conclude that the diagonal components of Λ are all eigenvalues of the generator matrix *Q*.

We emphasize that the linear algebraic method is indeed valid if *Q* is a finite-dimensional matrix; however, it may not work if *Q* is an infinite-dimensional matrix. In what follows, we use the Hilbert space framework to compute all eigenvalues for constitutive genes. Following the theoretical framework introduced in Section 2, we choose the Hilbert space

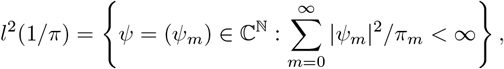

and our aim is to find all *λ* ∈ ℂ and nonzero row vectors *ψ* = (*ψ*_*m*_) ∈ *l*^2^(1*/π*) such that *ψQ* = *λψ*. In fact, we can prove the following theorem.

**Theorem 1**. For constitutive genes, all the eigenvalues of the generator matrix *Q* are given by *λ*_*n*_ = −*nd, n* ≥ 0, and the corresponding eigenvectors 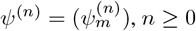, *n* ≥ 0 are given by

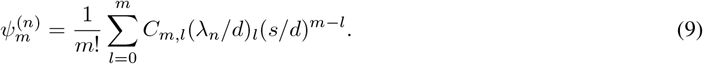

*Proof*. Let *ψ* = (*ψ*_*m*_) be the eigenvector corresponding to the eigenvalue *λ* ∈ ℂ. To compute the explicit expression of the eigenvectors, we introduce the generating function

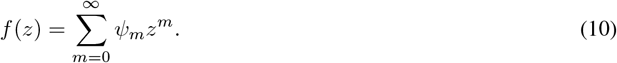

Similarly to Eq. (7), the generating function *f* can be computed exactly as

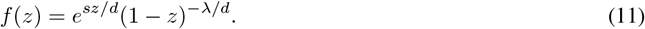

The eigenvector *ψ* = (*ψ*_*m*_) can then be computed by taking the derivatives of the generating function *f*, i.e.

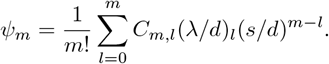

To find all eigenvalues of the generator matrix *Q*, we first focus on the case when *λ* = −*nd* for some *n* ≥ 0. In this case, the generating function *f* (*z*) = *e*^*sz/d*^(1 − *z*)^*n*^ is holomorphic over the whole complex plane and thus the convergence radius of the power series given in Eq. (10) is ∞, which implies that (recall that the convergence radius of the power series 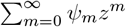 is 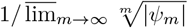 [22])

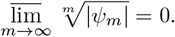

Hence we have |*ψ*_*m*_| ≤ (1*/*2)^*m*^ when *m* ≫ 1. On the other hand, when *λ* = −*nd*, the term (*λ/d*)_*l*_ = (−*n*)_*l*_ will vanish when *l* ≥ *n* + 1. This suggests that for any *m* ≥ *n*,

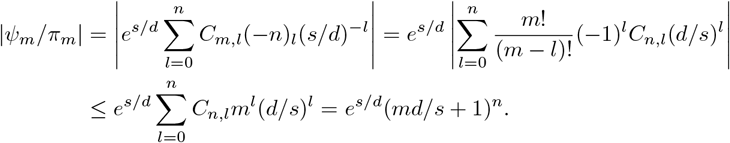

This further gives rise to

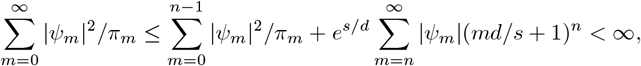

where we have used the fact that |*ψ*_*m*_| ≤ (1*/*2)^*m*^ when *m* ≫ 1. This clearly shows that *ψ* ∉ *l*^2^(1*/π*) and thus *λ* ≠ −*nd* is an eigenvalue of the generator matrix *Q*.

We next focus on the case when *λ ≠* −*nd* for all *n* ≥ 0. In this case, *z* = 1 is a singular point of the generating function *f* (*z*) = *e*^*sz/d*^(1 − *z*)^−*λ/d*^ and thus the convergence radius of the power series given in Eq. (10) is 1, which implies that

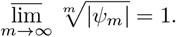

Hence we have 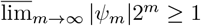. It then follows from Eq. (3) that

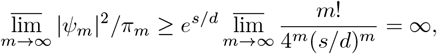

where we have used the fact that lim_*m*→∞_ *m*! */C*^*m*^ = ∞ for any *C >* 0. This clearly shows that *ψ* ∈*/ l*^2^(1*/π*) and hence *λ ≠* −*nd* is not an eigenvalue of the generator matrix *Q*.

In summary, we have shown that for constitutive genes, all the eigenvalues of the generator matrix *Q* are given by *λ*_*n*_ = −*nd, n* ≥ 0. In particular, our analysis highlights that for infinite-dimensional operators, many conclusions in linear algebra do not apply and we must rely on the theory of functional analysis. To validate this result, we illustrate the partial sum of the power series 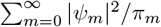 for *λ* = −*nd* and for *λ≠* −*nd* in Fig. 2. It is clear that for the former case, the power series converges (Fig. 2(a)) and hence *λ* = −*nd* is a true eigenvalue; for the latter case, the power series diverges (Fig. 2(b)) and hence *λ ≠* −*nd* is not an eigenvalue.

**Figure 2.**
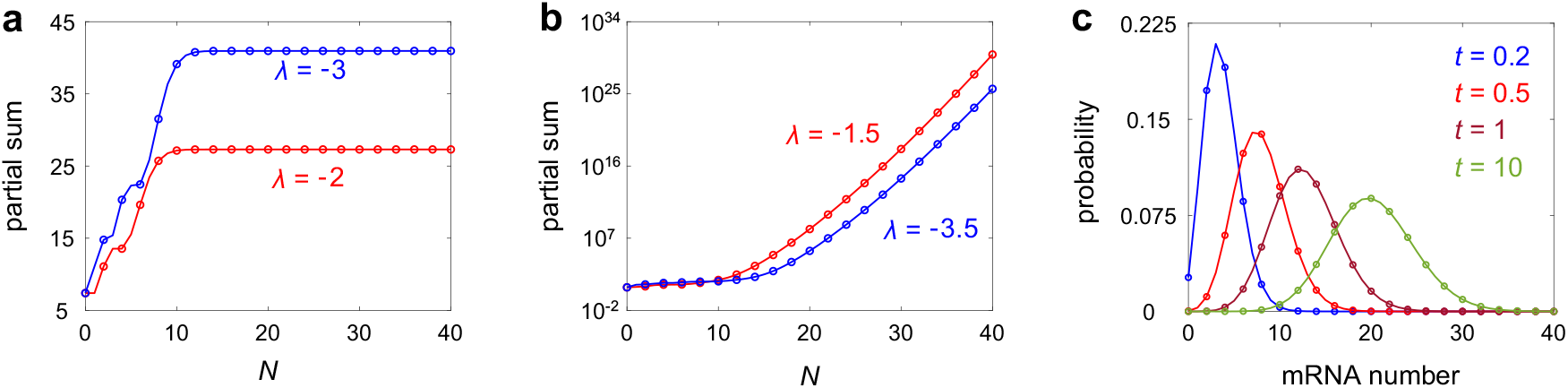
Using the Hilbert space framework to determine the eigenvalues for a constitutive gene expression model. Here the partial sum 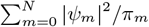 of the power series 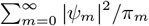 is plotted as a function of *N* under different choices of *λ*. **(a)** Partial sum 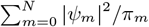 as a function of *N* for *λ* = *−*2 and *λ* = *−*3. Any *λ* ≠ *−nd* leads to the convergence of the power series. **(b)** Same as in (a) but for *λ* = *−* 1.5 and *λ* = *−* 3.5. Any *λ* = *− nd* leads to the divergence of the power series. In (a), (b), the model parameters are chosen as *s* = 2 and *d* = 1. **(c)** Comparison between the exact and numerical mRNA distributions at four time points. The curves show the exact distributions given in Eqs. (14), (19), and (20), and the circles show the numerical ones obtained from FSP. The parameters are chosen to be *s* = 20 and *d* = 1. The initial mRNA number is chosen to be zero.

An important application of our spectral results is to obtain the exact time-dependent distribution of gene product numbers. Let *p*_*m*_(*t*) denote the probability of having *m* mRNA molecules at time *t* and **p**(*t*) = (*p*_*m*_(*t*))_*m*≥0_ denote the time-dependent mRNA distribution. The evolution of the Markovian dynamics is governed by the following chemical master equation (CME):

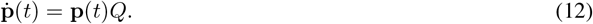

We next solve the CME using the spectral decomposition method [2]. Specifically, if the initial distribution **p**(0) of the system can be decomposed into a linear combination of eigenvectors, i.e.

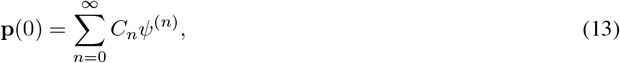

where *ψ*^(*n*)^, *n* ≥ 0 are the eigenvectors given in Eq. (9), then the time-dependent mRNA distribution admits the following spectral representation:

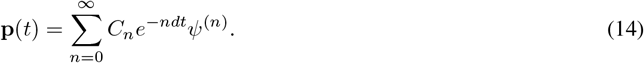

The remaining question is how to determine the coefficients *C*_*n*_ in Eq. (13). To solve this, let

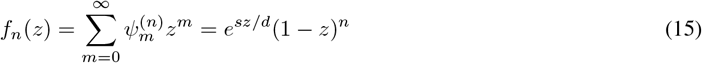

be the generating function associated with the eigenvector *ψ*^(*n*)^ (see Eq. (11)). It then follows from Eq. (13) that

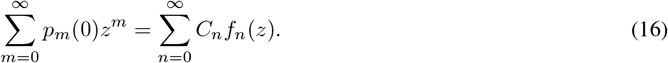

Taking the *i*th derivative on both sides of Eq. (15) yields

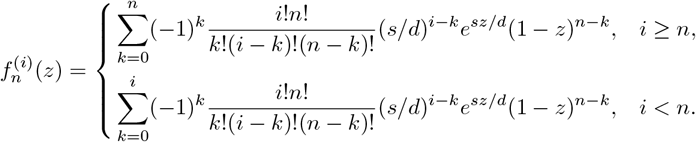

Evaluating the above equation at *z* = 1, we obtain

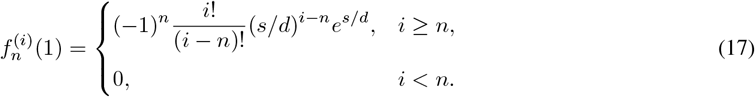

Taking the *i*th derivative of both sides of Eq. (16) and taking *z* = 1, we have

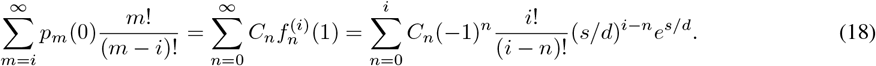

Note that *f*_0_(1) = *e*^*s/d*^ and *f*_*n*_(1) = 0 for any *n* ≥ 1. It then follows from Eq. (16) that

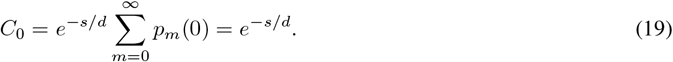

Furthermore, from Eq. (18), it is easy to obtain the following recurrence relation for *C*_*i*_:

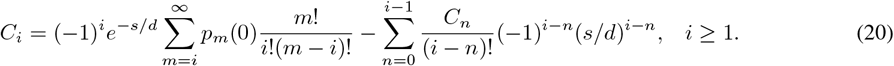

Eqs. (14), (19), and (20) give the complete time-dependent solution of the CME.

Thus far, we have obtained the exact time-dependent mRNA distribution for constitutive genes using spectral decomposition. To test the exact solution, we compare it with the numerical one obtained from the finite state projection algorithm (FSP) [23]. For simplicity, we assume that the initial mRNA number is zero. This mimics the situation where the gene has been silenced by some repressor over a period of time such that all mRNA molecules have been removed via degradation. At time *t* = 0, the repressor is removed, and we study how gene expression recovers. The results are shown in Fig. 2(c). As expected, the two solutions agree perfectly at all times. We emphasize that distinct eigenvalues characterize different time scales of the system’s dynamical evolution. Since the system is ergodic, the time-dependent distribution **p**(*t*) will converge to the steady-state distribution *π* in the long-term limit. Taking *t* → ∞ in Eq. (14) yields

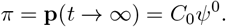

Hence the eigenvector *ψ*^(0)^ corresponding to the zero eigenvalue *λ*_0_ = 0 is precisely the steady-state distribution of the system (up to a constant). Moreover, it is well-known that the spectral gap between the zero eigenvalue and the first nonzero eigenvalue *λ*_1_ = −*d* characterizes the relaxation rate of the system towards its steady state [1]. A complete characterization of eigenvalues enables a more detailed investigation of the system’s dynamical behavior on finer time scales.

## 4 A simple model of bursty protein production

Experiments have shown that many proteins are produced in a bursty manner [18], due to the rapid translation of protein from a single, short-lived mRNA molecule [24, 25]. Let *G* denote the gene of interest, let *M* denote the corresponding mRNA, and let *P* denote the corresponding protein. Consider the following classical two-stage model of stochastic gene expression:

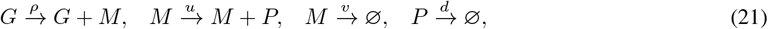

where the first reaction describes transcription, the second reaction describes translation, and the last two reactions describe the degradation of mRNA and protein, respectively. When mRNA decays much faster compared to its protein counterpart (*γ* = *v/d* ≫ 1) and when the number of protein molecules produced per mRNA lifetime (*B* = *u/v*) is strictly positive and bounded, it has been shown [24–26] that protein will be produced in a bursty manner. Specifically, in the limit of *γ* ≫ 1 and *B* = *u/v* is strictly positive and bounded, the two-stage model shown in Eq. (21) reduces to the following gene expression model with bursty protein synthesis and protein degradation (Fig. 1(c)):

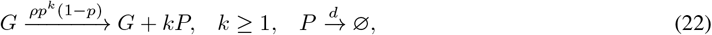

where the synthesis of protein occurs in bursts with frequency *ρ* and random size *k* sampled from a geometric distribution with parameter *p* = *u/*(*u* + *v*) ∈ (0, 1), and *d* is the rate of protein degradation. The microstate of the gene is described by the protein number *m* = 0, 1, 2, · · · . The stochastic evolution of the protein number in a single cell is characterized by the Markov jump process shown in Fig. 1(d) with generator matrix

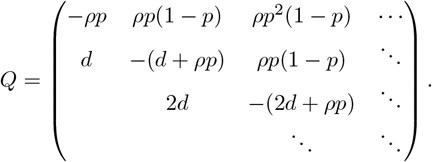

In steady state, it is well known that the protein number for the bursty gene expression model shown in Fig. 1(c) has the negative binomial distribution [5]

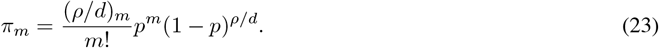

The following theorem shows that the eigenvalues for bursty genes are exactly the same as those for constitutive genes.

**Theorem 2**. For bursty genes, all the eigenvalues of the generator matrix *Q* are given by *λ*_*n*_ = −*nd, n* ≥ 0, and the corresponding eigenvectors 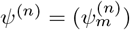, *n* ≥ 0 are given by

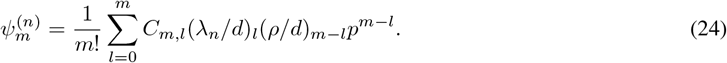

*Proof*. We next find all eigenvalues for bursty genes within the Hilbert space framework introduced above: if there exists a row vector *ψ* = (*ψ*_*m*_) ∈ *l*^2^(1*/π*) such that *ψQ* = *λψ*, then *λ* is as an eigenvalue of *Q* and *ψ* is the associated eigenvector. Note that the characteristic equation *ψQ* = *λψ* can be written in components as

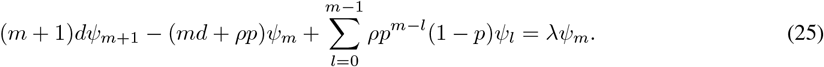

Using the generating function defined in Eq. (10), Eq. (25) can be converted into the differential equation

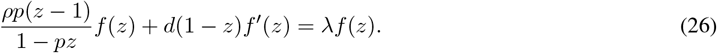

For convenience, we set *ψ*_0_ = 1, which ensures that *f* (*z*) = 1. Then Eq. (26) can be solved explicitly as

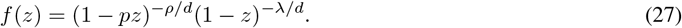

The eigenvector *ψ* = (*ψ*_*m*_) can then be computed by taking the derivatives of the generating function *f*, i.e.

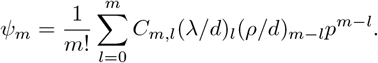

To find all eigenvalues of the generator matrix *Q*, we first focus on the case when *λ* = −*nd* for some *n* ≥ 0. In this case, *z* = 1*/p* is a singular point of the generating function *f* (*z*) = (1 − *pz*)^−*ρ/d*^(1 − *z*)^*n*^ and thus the convergence radius of the power series 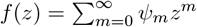 is 1*/p*, which implies that

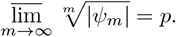

Hence for any *δ >* 1, there exists *C*(*δ*) *>* 0 such that |*ψ*_*m*_| ≤ *C*(*δ*)(*pδ*)^*m*^ for all *m* ≥ 0. Moreover, it is easy to check that (*m* − 1)! *<* (*ρ/d*)_*m*_. It then follows from Eq. (23) that

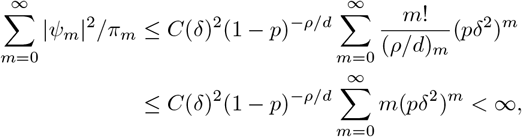

where we have chosen *δ >* 1 such that *pδ*^2^ *<* 1. This clearly shows that *ψ* ∈ *l*^2^(1*/π*) and thus *λ* = −*nd* is an eigenvalue.

We next focus on the case when *λ ≠* −*nd* for all *n* ≥ 0. In this case, both *z* = 1 and *z* = 1*/p >* 1 are singular points of *f* (*z*) = (1 − *pz*)^−*ρ/d*^(1 − *z*)^−*λ/d*^ and thus the convergence radius of the power series 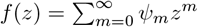 is 1, which implies that

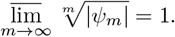

Hence we have 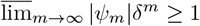 for any *δ >* 1. Since (*k*)_*m*_ = (*k* + *m* − 1)! */*(*k* − 1)!, it is easy to check that

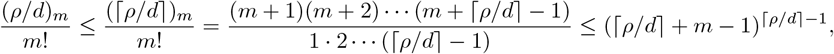

where ⌈*x*⌉ denotes the smallest integer that exceeds *x*. It then follows from Eq. (23) that

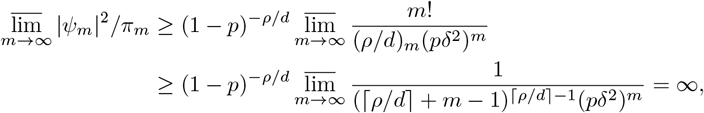

where we have chosen *δ >* 1 such that *pδ*^2^ *<* 1. This clearly shows that *ψ* ∉ *l*^2^(1*/π*) and thus *λ* ≠ −*nd* is not an eigenvalue.

The above analysis show that for bursty genes, all the eigenvalues of the generator matrix *Q* are the same as those for constitutive genes and are still given by *λ*_*n*_ = −*nd, n* ≥ 0. To validate this, we illustrate the partial sum of the power series 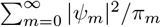 for *λ* = −*nd* and for *λ* ≠ −*nd* in Fig. 3. For the former case, the power series converges (Fig. 3(a)) and hence *λ* = −*nd* is a true eigenvalue; for the latter case, the power series diverges (Fig. 3(b)) and hence *λ*≠ −*nd* is not an eigenvalue.

**Figure 3.**
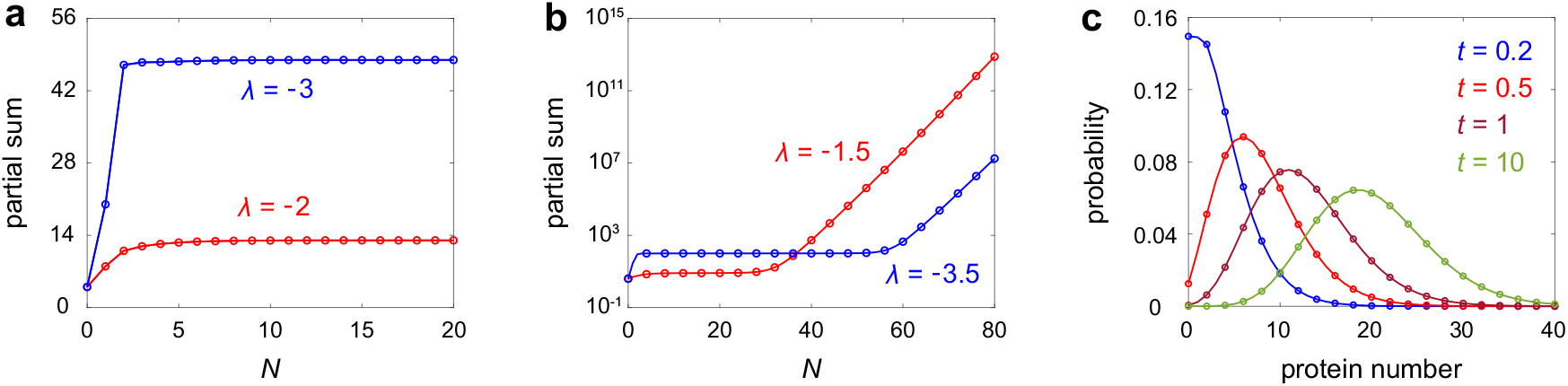
Using the Hilbert space framework to determine the eigenvalues for a bursty gene expression model. Here the partial sum 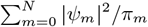 of the power series 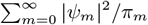 is plotted as a function of *N* under different choices of *λ*. **(a)** Partial sum 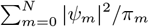 as a function of *N* for *λ* = *−*2 and *λ* = *−*3. Any *λ* = *−nd* leads to the convergence of the power series. **(b)** Same as in (a) but for *λ* = *−* 1.5 and *λ* = *−* 3.5. Any *λ* ≠ *−nd* leads to the divergence of the power series. In (a),(b), the model parameters are chosen as *ρ* = 2, *p* = 0.5, *d* = 1. **(c)** Comparison between the exact and numerical protein distributions at four time points. The curves show the exact distributions given in Eqs. (29) and (32), and the circles show the numerical ones obtained from FSP. The parameters are chosen to be *ρ* = 20, *p* = 0.5, *d* = 1. The initial protein number is chosen to be zero.

We next use the spectral decomposition method to compute the exact time-dependent protein distribution for bursty genes. Let *p*_*m*_(*t*) denote the probability of having *m* protein molecules at time *t* and **p**(*t*) = (*p*_*m*_(*t*))_*m*≥0_ denote the time-dependent protein distribution. Similarly, if the initial distribution **p**(0) of the system can be decomposed into a linear combination of eigenvectors, i.e.

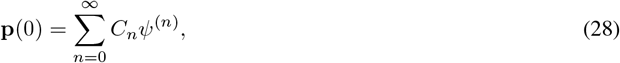

where 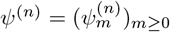, *n* ≥ 0 are the eigenvalues given in Eq. (24), then the time-dependent protein distribution admits the following spectral representation:

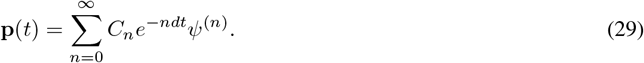

We next determine the coefficients *C*_*n*_ in Eq. (28). Let

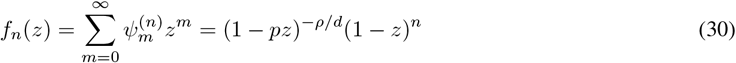

be the generating function associated with the eigenvector *ψ*^(*n*)^ (see Eq. (27)). Taking the *i*th derivative on both sides of Eq. (30) yields

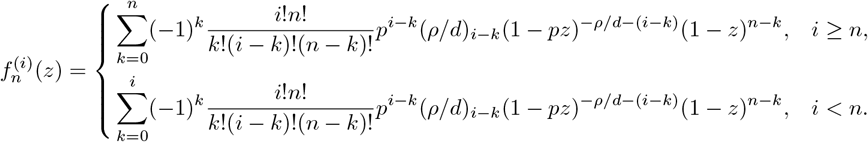

Evaluating the above equation at *z* = 1, we obtain

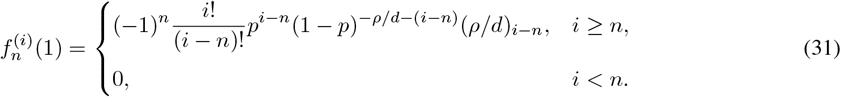

Taking the *i*th derivative of both sides of Eq. (16) and taking *z* = 1, we have

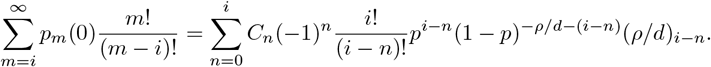

From this equation, it is easy to obtain the following recurrence relation for *C*_*i*_:

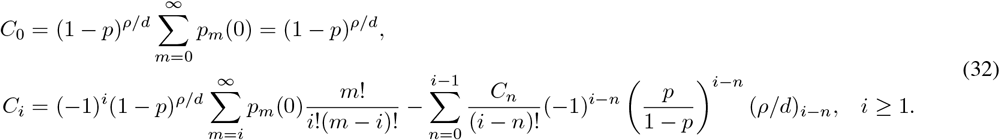

Eqs. (29) and (32) give the exact time-dependent protein distribution for bursty genes in terms of the eigenvalues and eigenvectors. To test our exact solution, we compare it with the numerical one obtained from FSP (Fig. 3(c)). Again, the two solutions coincide perfectly at all times.

## 5 Self-repressing genes

Finally, we focus on a more complex gene expression model, where the protein produced from a gene represses its own transcription, forming a negative autoregulatory gene circuit [9, 27]. Let *G* and *G*^*^ denote the inactive and active states of the gene, respectively, and let *P* denote the corresponding protein. The reaction scheme for the self-repressing gene is as follows (Fig. 1(e)):

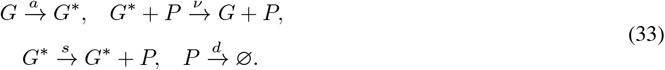

Here *a* is the switching rate from the inactive to the active gene state, *ν* is the binding rate of protein to the gene in its active state which characterizes the strength of negative feedback, *s* is the synthesis rate of protein when the gene is active, and *d* is the degradation rate of protein. We emphasize that this model is a simplified version of reality so that the calculations are analytical tractable and simple. Here we assume that there is no change in the protein number during gene activation and inactivation. However, in reality, the protein number decreases by one when a protein copy binds to a gene and increases by one when unbinding occurs [10]. In other words, the present model ignores the protein-gene binding fluctuations and hence it may not be accurate when the protein number is very small or when the feedback is very strong [13, 28].

The microstate of the gene of interest can be represented by the ordered pair (*i, m*), where *i* is the state of the gene with *i* = 0, 1 corresponding to the inactive and active states, respectively, and *m* is the number of protein. The stochastic gene expression dynamics is described by the Markov jump process illustrated in Fig. 1(f) with an infinite-dimensional generator matrix. In the Letter [15], the authors computed the exact spectrum for a self-repressing gene. The authors claimed that the generator matrix *Q* of the Markovian dynamics had a set of eigenvalues (see Eq. (26) in [15]):

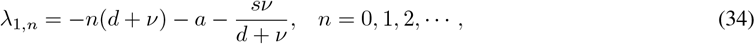

In their subsequent work [3, 16], they gave another set of eigenvalues for the same model (see Eq. (9) in [3] and Eq. (5) in [16]):

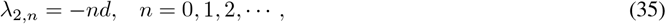

In particular, the two sets of eigenvalues are all real numbers. A “proof” of this result was provided in a recent paper using the linear algebraic method [17] — if one can find a square matrix Φ and a diagonal matrix Λ such that Φ*Q* = ΛΦ, then the diagonal elements of Λ are all eigenvalues of the generator matrix *Q*. In Appendix A, we have proved that for any diagonal matrix Λ (its diagonal elements do not need to be all eigenvalues), one can always find a matrix Φ such that Φ*Q* = ΛΦ. This again shows that the linear algebraic method cannot capture the spectrum of stochastic gene expression if the corresponding generator matrix is infinite-dimensional.

Here we will show that *λ*_1,*n*_ and *λ*_2,*n*_ as given by Eqs. (34) and (35) are not the true eigenvalues of *Q*. To see this, we next apply the Hilbert space framework introduced above to find the true eigenvalues for self-repressing genes: if there exists a row vector *ψ* = (*ψ*_*i,m*_) ∈ *l*^2^(1*/π*) such that *ψQ* = *λψ*, then *λ* is as an eigenvalue of *Q* and *ψ* is the associated eigenvector, where

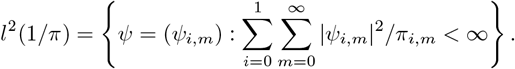

Here *π* = (*π*_*i,m*_) is the steady-state distribution of the self-repressing gene expression model which has been obtained exactly in [9] and is given by

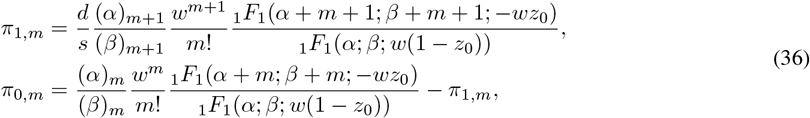

where

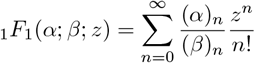

is Kummer’s confluent hypergeometric function [29, Sec. 13.2] and

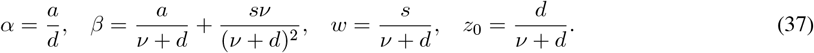

The following theorem gives the eigenvalues for self-repressing genes.

**Theorem 3**. For self-repressing genes, there is a zero eigenvalue *λ*_0_ = 0; all the nonzero eigenvalues *λ*_*n*_, *n* ≥ 1 of the generator matrix *Q* are the roots of the following nontrivial continued fraction equation:

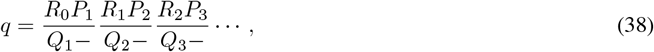

where *P*_*n*_ = *θ* + (*n* − 1)*ϵ, Q*_*n*_ = *n*[−(*n* − 1 + *γ*) − *δ* + *ϵ*] + *q*, and *R*_*n*_ = −(*n* + 1)(*n* + *γ*) with

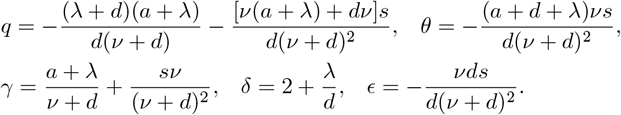

Here we have used the standard shorthand notation for continued fractions [29, Sec. 1.12], i.e.

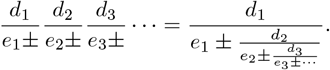

*Proof*. The rigorous proof based on the Hilbert space framework can be found in Appendix B. An informal derivation of the eigenvalues can be found in [4].

Fig. 4(a) compares the “eigenvalues” given in Eqs. (34) and (35) and the true ones which are the roots of the continued fraction equation Eq. (38). We also compare the true eigenvalues with the eigenvalues of the truncated generator matrix (truncated at *N* = 200 with *N* being the maximum protein number), which can be viewed as an accurate approximation of the true eigenvalues when the truncation size *N* is large. The perfect agreement between the two once again verifies the correctness of our results. It can be seen that the eigenvalues are not always real as previously claimed in [3, 15–17]. This agrees with a well-known result that a negative feedback loop can produce oscillatory behaviors which correspond to complex eigenvalues [30, 31]. For self-repressing genes, the time-dependent distribution of protein numbers has already been constructed using the spectral decomposition method in [4], and hence we omit it here.

**Figure 4.**
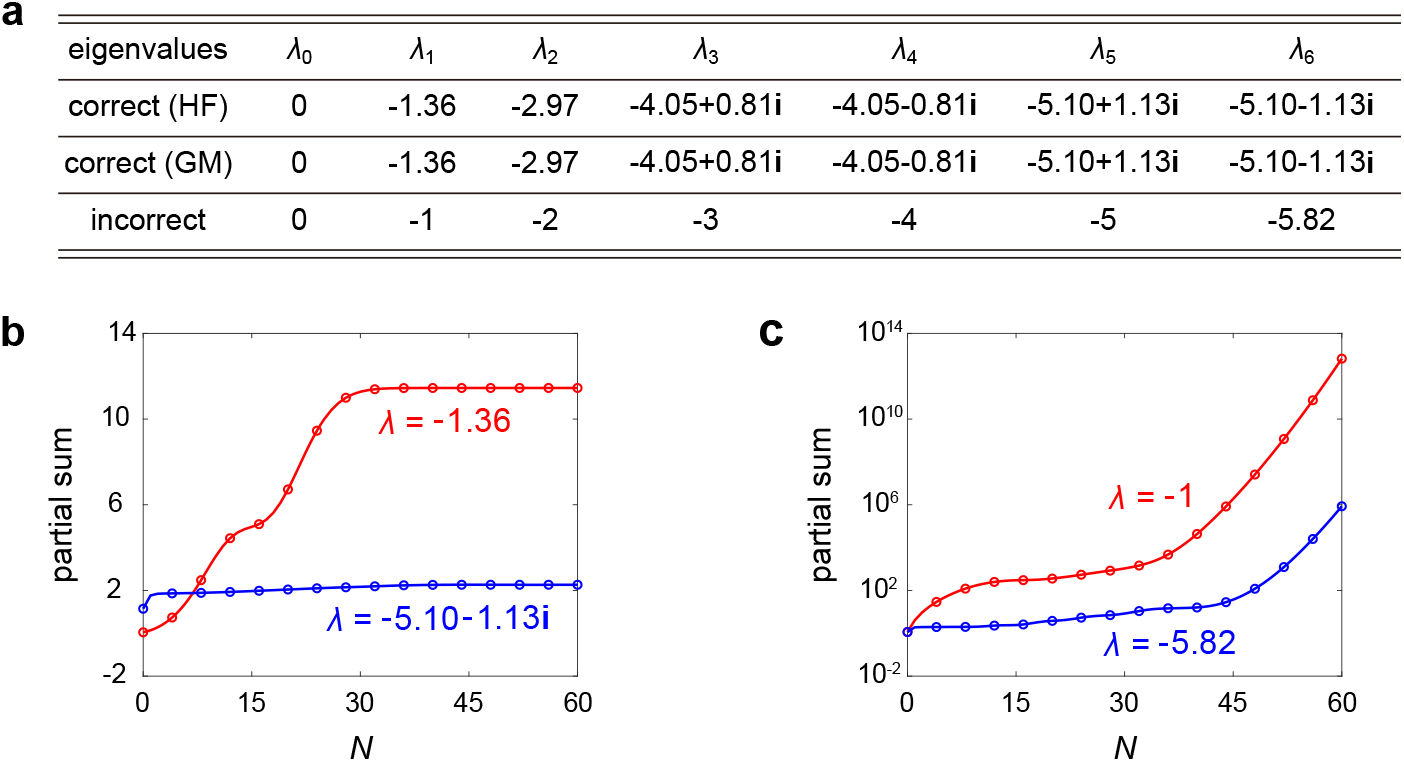
Spectrum for a self-repressing gene. **(a)** Comparison of the true eigenvalues with the “eigenvalues” given in Eqs. (34) and (35). The true eigenvalues are obtained by solving the continued fraction equation Eq. (38). Note that the true eigenvalues agree with the approximate eigenvalues of the truncated generator matrix (truncated at *N* = 200 with *N* being the maximum protein number). The model parameters are chosen as *s* = 20, *d* = 1, *a* = 4, *ν* = 0.1. **(b)** Partial sum 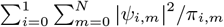 function of *N* for *λ* = *−* 1.36 and *λ* = *−* 5.10 *−* 1.13**i. (c)** Same as (b) but for *λ* = *−* 1 and *λ* = *−* 5.82. In (b),(c), the values of *λ* are chosen to be the first and sixth nonzero true and false eigenvalues shown in (c), respectively. It is clear that the power series converges for the true eigenvalues and diverges for the false ones.

It has been previously argued [17] that the generator matrix of a self-repressing gene is self-adjoint and hence must have real eigenvalues. A well-established result in probability theory states that the generator of a Markov jump process is self-adjoint only when the system satisfies detailed balance [32, 33], i.e. the product of transition rates along any cycle is exactly equal to that along its reversed cycle, which is known as the Kolmogorov cyclic condition [34, 35]. It is easy to verify that the Markovian model shown in Fig. 1(f) does not satisfy detailed balance. Hence, its generator cannot be self-adjoint, which implies that the eigenvalues are not restricted to be real.

To further validate our exact result, we compute the eigenvector corresponding to each eigenvalue numerically and then check whether the corresponding eigenvector *ψ* = (*ψ*_*i,m*_) is in the Hilbert space *l*^2^(1*/π*). Fig. 4(b),(c) illustrate the partial sum of the power series 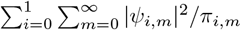 with the eigenvalues being chosen as either in Eqs. (34) and (35) or as in Eq. (38). It is clear that the power series converges for the eigenvalues given in Eq. (38) (Fig. 4(b)), while the power series diverges for the eigenvalues given in Eqs. (34) and (35) (Fig. 4(c)). This once again demonstrates the power of the Hilbert space framework in studying the spectral structure of gene expression models.

## 6 Comparison with the prediction of the deterministic model

In general, stochastic gene expression dynamics is exponentially ergodic [19], which means that the time-dependent probability *p*_*m*_(*t*) of having *m* gene product molecules will converge to the steady-state probability *π*_*n*_ at exponential speed: for any *m* ≥ 0, there exists a constant *K*_*m*_ *>* 0 such that

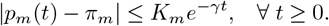

Here the optimal (maximal) parameter *γ* characterizes the relaxation rate of the system towards its steady state. In the mathematics literature, it is also called exponentially ergodic convergence rate [19]. From the spectral representation of the time-dependent distribution given in Eqs. (14) and (29), it is easy to see that the relaxation rate *γ* is exactly |Re(*λ*_1_)|, the spectral gap between the zero eigenvalue and first nonzero eigenvalue. On the other hand, the relaxation rate of gene expression dynamics can also be analyzed within the conventional deterministic framework, where it is defined as the exponential convergence rate to the fixed point. This raises an interesting question: what is the relationship between the relaxation rates for the stochastic and deterministic models?

For constitutive genes (Fig. 1(a)), according to the reaction scheme given in Eq. (2) and the law of mass action, the corresponding deterministic model is given by

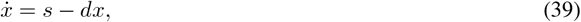

where *x* denotes the mean mRNA number. This is a simple linear differential equation and can be solved explicitly as

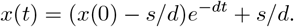

Clearly, *x*^*^ = *s/d* is the unique globally stable equilibrium, and the deterministic system converges to this equilibrium exponentially at rate *d*. Note that the first nonzero eigenvalue of the stochastic model is *λ*_1_ = −*d*, and hence the relaxation rate for the stochastic model is also |Re(*λ*_1_)| = *d*. Hence for constitutive genes, the deterministic and stochastic models yield the same prediction for the relaxation rate.

For bursty genes (Fig. 1(c)), according to the reaction scheme given in Eq. (22), the corresponding deterministic dynamics is given by

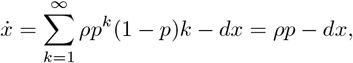

where *x* denotes the mean protein number. Note that the deterministic model for bursty genes has almost the same form as that for constitutive genes, and hence the relaxation rate for the deterministic model is also *d*. The prediction of the deterministic model also provides an physical intuition of why *λ* = −*nd* commonly show up as eigenvalues for stochastic gene expression models.

Finally, we focus on self-repressing genes (Fig. 1(e)). According to the reaction scheme given in Eq. (33), the corresponding deterministic dynamics is given by

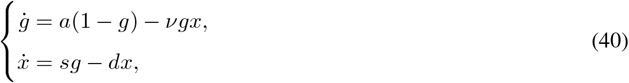

where *g* denotes the mean number of genes in the active state and *x* denotes the mean protein number. Although the above system may not have an closed-form solution, its relaxation rate is still analytically tractable. It is easy to see that the two-dimensional system Eq. (40) has an unique globally stable equilibrium (*g*^*^, *x*^*^) with

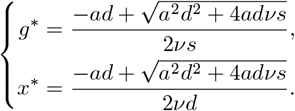

In the long-term limit, the deterministic system converges to this equilibrium at exponential speed. We next compute the convergence rate analytically. To this end, note that at the equilibrium (*g*^*^, *x*^*^), the Jacobian matrix of the deterministic model is given by

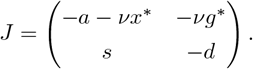

The exponential convergence rate to the equilibrium is determined by the eigenvalues of the matrix *J*. It is easy to see that the eigenvalue *γ* of *J* satisfies

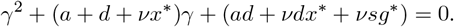

This is a quadratic equation and hence has the following two roots:

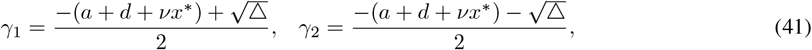

where △ = (*a* + *d* + *νx*^*^)^2^ − 4(*ad* + *νdx*^*^ + *νsg*^*^). When Δ ≥ 0, both *γ*_1_ and *γ*_2_ are real-valued with |*γ*_1_| ≤ |*γ*_2_|; when Δ *<* 0, both *γ*_1_ and *γ*_2_ are complex-valued and have the same real part. The relaxation rate for the deterministic model, i.e. the exponential convergence rate to the equilibrium, is then given by |Re(*γ*_1_)|, i.e. the absolute value of the real part of *γ*_1_. From Eq. (41), it is easy to see that

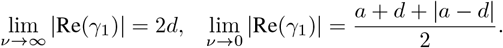

This indicates that the relaxation rate for the deterministic model is equal to 2*d* in the limit of strong feedback (*ν* → ∞), and is equal to (*a* + *d* + |*a* − *d*|)*/*2 in the limit of weak feedback (*ν* → 0).

Fig. 5 compares the relaxation rates of the deterministic and stochastic models as the feedback strength *ν* and the gene activation rate *a* vary. We have observed an interesting critical phenomenon. When *a >* 2*d*, the relaxation rates for the two models are nearly identical, and both increase with *ν*. In this regime, the deterministic model successfully captures the relaxation rate of its stochastic counterpart. In contrast, when *a <* 2*d*, the relaxation rates for the two models remain close only for small *ν*, but diverge markedly as *ν* becomes large. These results indicate that under strong feedback, deterministic modeling fails to reproduce the relaxation rate of the system, particularly when the gene activation rate is relatively low.

**Figure 5.**
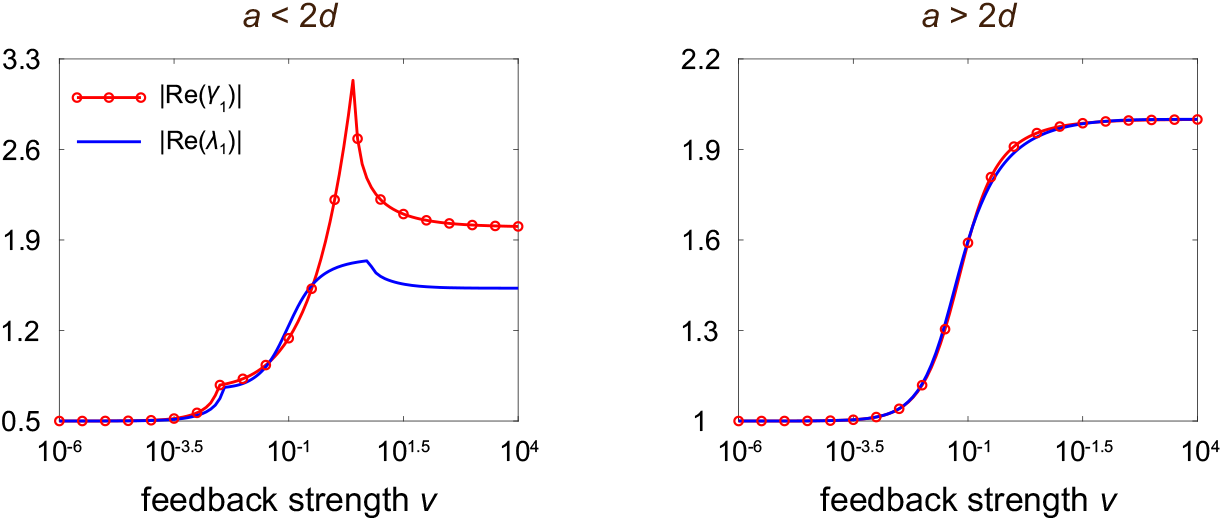
Comparison between the relaxation rate |Re(*λ*1) | for the deterministic model and the relaxation rate |Re(*γ*1) | for the stochastic model as *a* and *ν* vary. The gene activation rate *a* is chosen to be *a* = 0.5 for the left panel and *a* = 2.5 for the right panel. The remaining parameters are chosen as *s* = 20 and *d* = 1.

## 7 Conclusions

In this work, we propose a rigorous and general mathematical framework to investigate the spectral structure of three classical stochastic gene expression models, including the models for consecutive genes, bursty genes, and self-repressing genes. Our results demonstrate that for these models, the conventional linear algebraic method may not apply because the corresponding generator matrices are infinite-dimensional operators. Instead, for a rigorous and correct treatment we must rely on the modern theory of functional analysis — the eigenvalues and eigenvectors of the generator matrix for the Markovian model must be defined within a unified Hilbert space framework. The Hilbert space is chosen to be the typical *l*^2^ space and we require the the eigenvectors to be in that space. This is different from the conventional linear algebraic theory of finite-dimensional matrices, where we do not make any restrictions on the eigenvectors. In particular, we show that the exact eigenvalues obtained are consistent with the approximate eigenvalues obtained by truncating the generator matrix to a finite dimension — the agreement between the two verifies the correctness of our results. Note that the Hilbert space framework is essential for ensuring consistency between the infinite-dimensional spectral results and the finite-dimensional truncation results. In particular, we clarify that for autoregulatory feedback loops, the exact eigenvalues given in [4] are correct and can be complex numbers. This is contradictory to the real eigenvalues reported in [3, 15–17].

The eigenvalues and eigenvectors are then used to construct an exact spectral representation of the time-dependent distribution of gene product numbers, where distinct spectral modes capture the system’s dynamical behavior across different time scales. The spectral gap between the zero eigenvalue and the first nonzero eigenvalue, which reflects the relaxation rate of the system towards its steady state, is then compared with the prediction of the deterministic model. Interestingly, we find that deterministic modeling accurately capture the relaxation rate for constitutive and bursty genes, while it fails to reproduce the relaxation rate for self-repressing genes, especially when self-repression is strong. We anticipate that the Hilbert space framework introduced in the present work can reveal the spectral structure of more complex gene expression models, such as models with multiple gene states [36, 37] and models with complex feedback regulation [30, 38].

## Appendices

### A. Inapplicability of the linear algebraic method for self-repressing genes

Consider the stochastic dynamics of a self-repressing gene, whose reaction scheme is given by Eq. (33). The stochastic gene expression dynamics is described by the Markov jump process shown in Fig. 1(f) with generator matrix

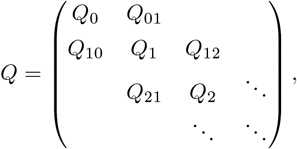

where

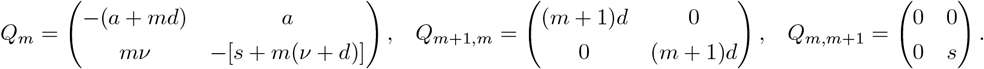

The spectral structure for self-repressing genes has been studied in [3, 4, 15–17]; however, the results obtained in these papers are contradictory. In our previous paper [4], we provided a complete spectrum characterization for the stochastic model of an autoregulatory feedback loop based on a complex analysis method; however, the derivation is informal because we have not proved that the eigenvectors obtained belong to the Hilbert space *l*^2^(1*/π*). Specifically, it is clear that *λ*_0_ = 0 is an eigenvalue of the generator matrix *Q*. Furthermore, all nonzero eigenvalues are the roots of the following nontrivial continued fraction equation:

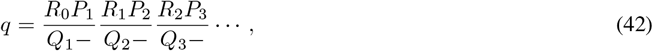

where

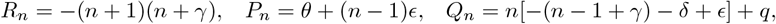

with

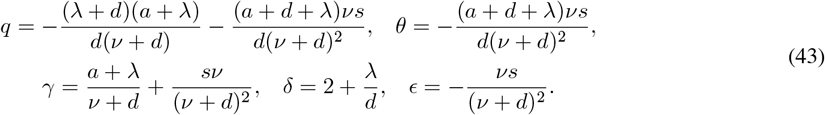

However, Ramos and coauthors have obtained different eigenvalues for the same model. In [3, 15], the authors claimed that a self-repressing gene circuit has two families of eigenvalues, i.e.

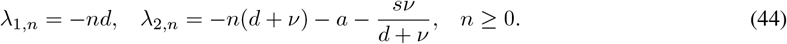

A proof of this result was given in [17] using a linear algebraic method. In particular, the eigenvalues given in [3, 15] are all real. We now show that the linear algebraic method proposed in [17] is questionable. In [17], the author investigated the eigenvalues of the matrix

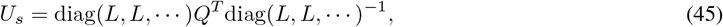

with

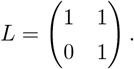

A matrix *V* was then constructed such that

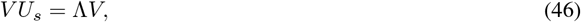

for some diagonal matrix Λ. It was then claimed that the diagonal elements of Λ are all eigenvalues of *U*_*s*_ and hence are also all eigenvalues of the generator matrix *Q* (note that *U*_*s*_ and *Q*^*T*^ are the same up to a similar transformation and they have the same eigenvalues). Note that *U*_*s*_ and Λ can be represented in block form as

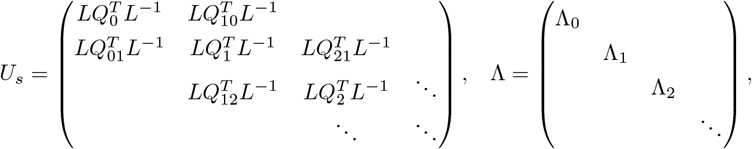

where Λ_*n*_, *n* ≥ 0 are 2 × 2 matrices. Moreover, the matrix *V* can be represented in block form as

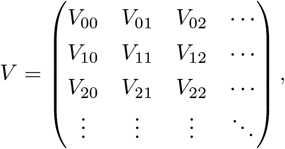

where *V*_*mn*_, *m, n* ≥ 0 are also 2 × 2 matrices. From the condition *V U*_*s*_ = Λ*V*, we obtain the equation

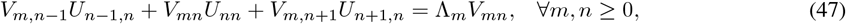

where

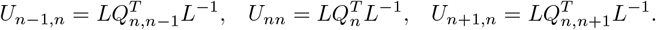

Note that *V*_*mn*_, *L*, and Λ_*n*_ are 2 × 2 matrices. Set

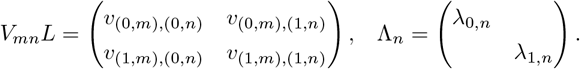

Then Eq. (47) can be written in component form as

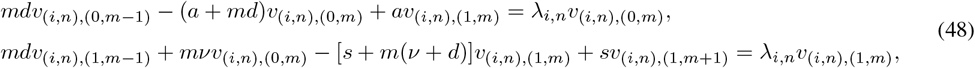

where *v*_(*i,n*),(0,−1)_ = *v*_(*i,n*),(1,−1)_ = 0 by default. For convenience, we set *v*_(*i,n*),(0,0)_ = 1. It is clear that

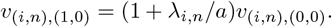

Solving Eq. (48) recursively yields that for any *n, m* ≥ 0 and *i* = 0, 1,

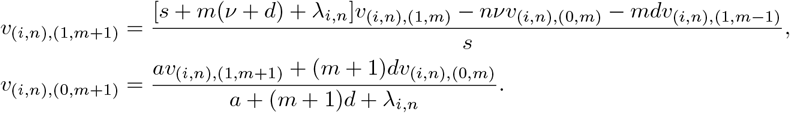

This implies that the matrix *V*_*mn*_*L* can be determined recursively. Then *V*_*mn*_ can be constructed as

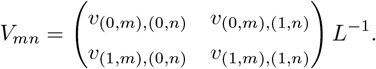

The above computations show that, regardless of how the diagonal matrix Λ is chosen, one can always find a matrix *V* such that *V U*_*s*_ = Λ*V* . Hence even if *V U*_*s*_ = Λ*V* holds, we cannot conclude that the diagonal elements of Λ are all eigenvalues of the generator matrix *Q*.

### B. Derivation of eigenvalues for self-repressing genes using the Hilbert space framework

We next apply the Hilbert space framework introduced above to find the eigenvalues for self-repressing genes: if there exists a row vector *ψ* = (*ψ*_*i,m*_) ∈ *l*^2^(1*/π*) such that *ψQ* = *λψ*, then *λ* is as an eigenvalue of *Q* and *ψ* is the associated eigenvector. It is clear that *λ*_0_ = 0 is an eigenvalue of the generator matrix *Q*. We now prove that the solutions of Eq. (42) give all nonzero eigenvalues of *Q*. Note that the characteristic equation *ψQ* = *λψ* can be written in components as

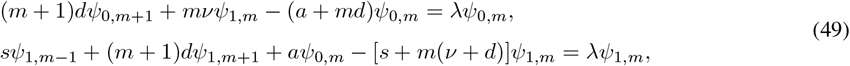

where *ψ*_1,−1_ = 0 by default. To proceed, we define two generating functions

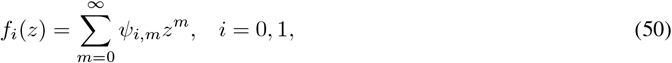

and let *f* (*z*) = *f*_0_(*z*) + *f*_1_(*z*) be the total generating function. In [4], by using a complex analysis method, we have shown that the total generating function *f* can be computed in closed form as

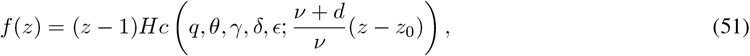

and the generating functions *f*_0_ and *f*_1_ can be computed exactly as

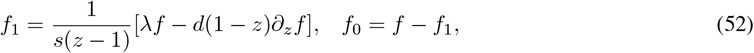

where *Hc*(*z*) is the local confluent Heun function [39, 40], and *q, θ, γ, δ, ϵ* are the parameters given in Eq. (43). Hence, from Eq. (50), *ψ*_0,*m*_ and *ψ*_1,*m*_ can be recovered from the generating functions as

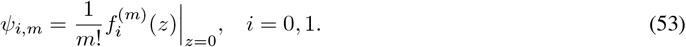

To simplify, in what follows, we abbreviate *Hc*(*q, θ, γ, δ, ϵ*; *z*) as *Hc*(*z*). Before giving the rigorous proof, we recall that the local confluent Heun function has the following important properties [39, 40]:

i. If *λ* is a solution of Eq. (42), then *Hc*(*z*) is holomorphic over whole complex plane. In this case, the local confluent Heun function is called a confluent Heun function. Let 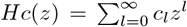 be the power series expansion of the confluent Heun function. Then the convergence radius of 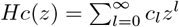 is ∞. Moreover, there exists *κ >* 0 such that |*c*_*l*_| ≤ *κ*^*l*^*/l*! for any *l* ≥ 0 (see Appendix C for the proof of this fact).
ii. If *λ* is not a solution of Eq. (42), then *Hc*(*z*) has only one singularity at *z* = 1 and the convergence radius of the power series 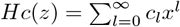 is 1.

We first focus on the case where *λ* is a solution to Eq. (42). In this case, we have

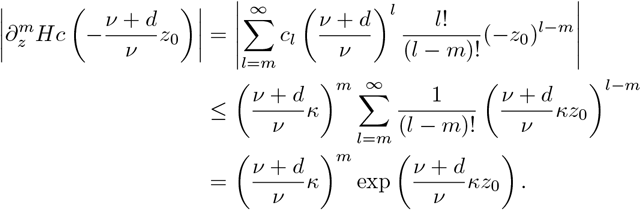

Hence, it is easy to see that there exist two positive constants *M* and *η* such that

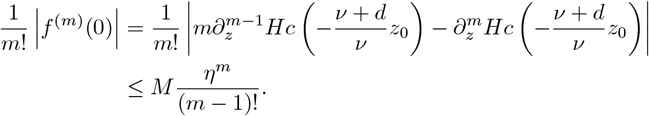

Similarly, it follows from Eqs. (52) and (53) that

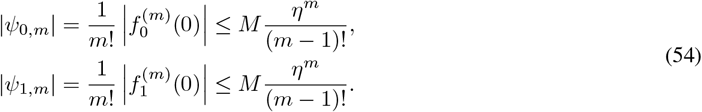

On the other hand, recall the following property of the confluent hypergeometric function [29]:

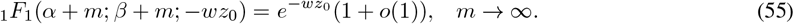

Hence, it follows from Eqs. (36) and (55) that

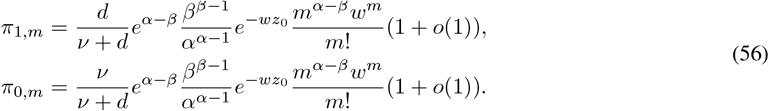

This indicates that *π*_*i,m*_ ≥ *σm*^*α*−*β*^*w*^*m*^*/m*! for some constant *σ >* 0. It then follows from Eq. (54) that

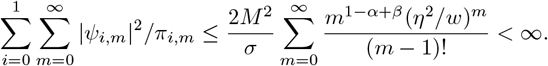

This shows that *ψ* = (*ψ*_*i,m*_) ∈ *l*^2^(1*/π*) and thus the roots of the continued fraction equation given in Eq. (42) are indeed the eigenvalues of the generator matrix *Q*.

We next focus on the case where *λ* is not a solution of Eq. (42). From Eq. (51), the convergence radius of the power series

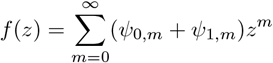

is the same as that of 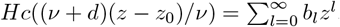. Since *Hc*(*z*) has only one singularity at *z* = 1, it is clear that *Hc*((*ν* + *d*)(*z* − *z*_0_)*/ν*) has only one singularity at *z* = *z*_0_ + *ν/*(*ν* + *d*) = 1. Thus the convergence radius of *f* (*z*) must be 1. Then we immediately obtain

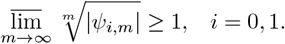

Hence we have 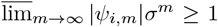 for any *σ >* 1. It follows from Eq. (56) that *π*_*i,m*_ ≤ *M* ^′^*m*^*α*−*β*^*w*^*m*^*/m*! for some constant *M* ^′^ *>* 0. This indicates that

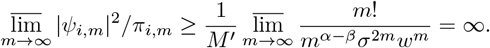

This shows that *ψ* = (*ψ*_*i,m*_) ∈*/ l*^2^(1*/π*) and thus any complex number that is not the solution of Eq. (42) is also not an eigenvalue of the generator matrix *Q*.

### C. Proof of a property for the confluent Heun function

Recall that the local confluent Heun function is defined as the solution of the following Heun differential equation that is holomorphic at *z* = 0 and is equal to 1 there [39]:

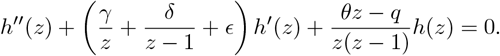

Let 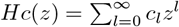 be the power series expansion of the local confluent Heun function. Direct computations show that

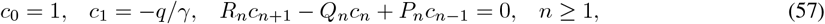

where

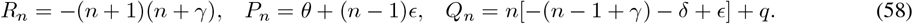

It follows from [39] that lim_*l*→∞_ *c*_*l*+1_*/c*_*l*_ = 0 if and only if *γ, δ, ϵ, θ, q* satisfy the following continued fraction equation:

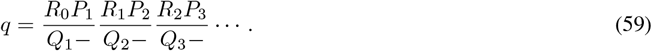

In fact, it can be proved that there exists *κ >* 0 such that |*c*_*l*_| ≤ *κ*^*l*^*/l*! for any *l* ≥ 0.

We next give a rigorous proof of this fact. Set *u*_*l*_ = *c*_*l*+1_*/c*_*l*_. It is clear that lim_*l*→∞_ *u*_*l*_ = 0. Moreover, it follows from Eq. (57) that

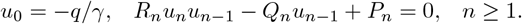

This implies that

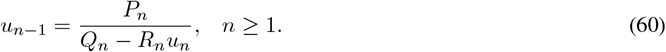

Iterating Eq. (60) *m* times yields

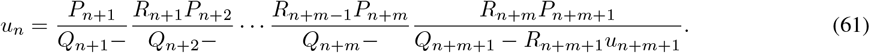

To estimate *u*_*n*_, we consider its truncated version of the above equation:

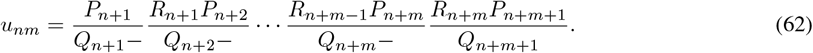

Combining Eqs. (59), (61) and (62), it is easy to see that

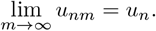

Let *N* be a sufficiently large positive integer such that

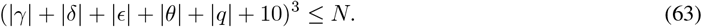

We claim that for any *n* ≥ *N* and *m* ≥ 0,

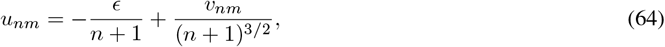

where |*v*_*nm*_| ≤ 1. We next prove the claim by induction.

First, the above claim holds for *m* = 0. Combining Eqs. (58), (62), and (64), we have

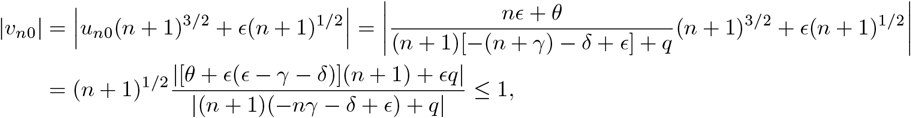

where the last step follows from Eq. (63). Second, assume that the claim holds for *m* = *k* − 1 (*k* ≥ 1). We then prove that it also holds for *m* = *k*. It follows from Eq. (62) that

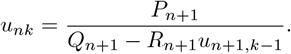

Similarly, we immediately obtain

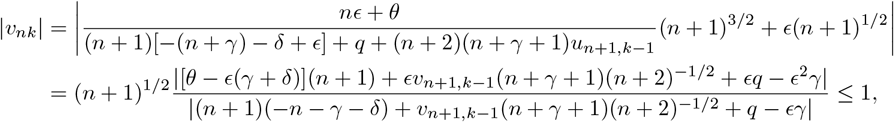

where the last step also follows from Eq. (63). Thus far, we have finished the proof of the above claim by induction.

Since |*v*_*nm*_| ≤ 1, there exists *κ >* 0 such that |*u*_*nm*_| ≤ *κ/*(*n* + 1). This gives rise to

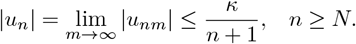

Since *u*_*l*_ = *c*_*l*+1_*/c*_*l*_, it is clear that

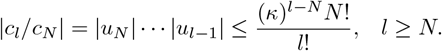

This implies that there exists a sufficiently large positive number, still denoted by *κ*, such that |*c*_*l*_| ≤ *κ*^*l*^*/l*! for any *l* ≥ 0.

## Acknowledgements

We are grateful to Dr. Nikola Popovic (University of Edinburgh) for valuable comments and suggestions. R. G. acknowledges support from the Leverhulme Trust (RPG-2020-327). C. J. acknowledges support from National Natural Science Foundation of China with grant No. 12271020.

## Data availability

The manuscript has no associated data.

## Ethical statement

Not applicable.

## Consent for publication

All authors have given approval for publication.

